# Lentiviral co-packaging mitigates the effects of intermolecular recombination and multiple integrations in pooled genetic screens

**DOI:** 10.1101/262121

**Authors:** David Feldman, Avtar Singh, Anthony J. Garrity, Paul C. Blainey

## Abstract

Lentiviral vectors are widely used for functional genomic screens, enabling efficient and stable transduction of target cells with libraries of genetic elements. Unfortunately, designs that rely on integrating multiple variable sequences, such as combinatorial perturbations or perturbations linked to barcodes, may be compromised by unintended consequences of lentiviral packaging. Intermolecular recombination between library elements and integration of multiple perturbations (even at limiting virus dilution) can negatively impact the sensitivity of pooled screens. Here, we describe a simple approach to prevent recombination between lentiviral vectors containing multiple linked variable elements, such as the recently reported CRISP-seq, Perturb-seq, and Mosaic-seq designs. We show that modifying the packaging protocol by diluting the perturbation library with a carrier plasmid increases the fraction of correct, single integrations from <60% to >90%, at the cost of reducing titer by 100-fold.

## Introduction

Lentiviral vectors provide a convenient, scalable platform to deliver genetic perturbations to cells *en masse* and read out the identity of each perturbation by next-generation sequencing^1,2^. There is increasing interest in screening approaches reliant on the delivery of multiple library elements to each cell, for example, in CRISPR-based single cell gene expression screens^3–8^. Such approaches facilitate the study of genetic interactions by probing cells with combinations of perturbations or convenient detection of perturbations by readout of a barcode sequence. However, the goal of accurately delivering a single integrated library variant per cell is complicated by aspects of lentiviral delivery.

Lentiviral virions normally contain two copies of the viral genome. During standard lentivirus production, transfection of packaging cells with multiple plasmids generates virions containing two distinct library elements, which can then lead to intermolecular recombination that shuffles variable library sequences (“barcode swapping”). Together with the inadvertent integration of multiple variants in individual target cells, this process has the effect of reducing the sensitivity of pooled screens. For screens where all library elements are read out (e.g. targeted pairs of gene knockouts), recombination events can be detected and filtered out before statistical analysis^6,9^. However, in situations where functional library elements are not sequenced directly, but are rather inferred via a linked barcode, recombination or multiple integration can lead to mislabeled data and has been noted to decrease the statistical power of genetic screens at a given number of cells analyzed^10,11^.

Recombination can arise from the template-switching activity of the lentiviral reverse-transcriptase^12^. As the lentivirus capsid normally packages a dimer of RNA genomes, intermolecular recombination could in principle occur in target cells infected by a single virion. The fraction of target cells with recombined integrants depends on the distance between variable sequences and has been measured to exceed 30% for distances greater than 1 kb^13^. Such wide spacing of library elements is common when the elements are separated by regulatory sequences or when an element is used as a 3’ barcode in an expressed transcript^10,11,13^.

To quantify the frequencies of barcode swapping and multiple integration events, we performed clonal analysis of target cells transduced with a library of CRISPR sgRNAs and transcribed barcode elements separated by >1.7 kb. Similar to other groups, we found that standard lentiviral packaging results in substantial (>30%) barcode swapping between library elements. We further observed that an unexpectedly high number of target cells had multiple library variants integrated into their genomes even when transduced at low multiplicity-of-infection (MOI). Here we show that by diluting the perturbation library plasmid with a sufficient excess of carrier plasmid during the packaging step (Figure 1), we were able to substantially reduce barcode swaps (<4%) and attenuate the rate of multiple integrations several-fold. Altogether, this co-packaging strategy constitutes a simple solution to improve data quality for genetic screens without constraining library vector design or necessitating individual (“arrayed”) packaging of library elements.

**Figure 1:**
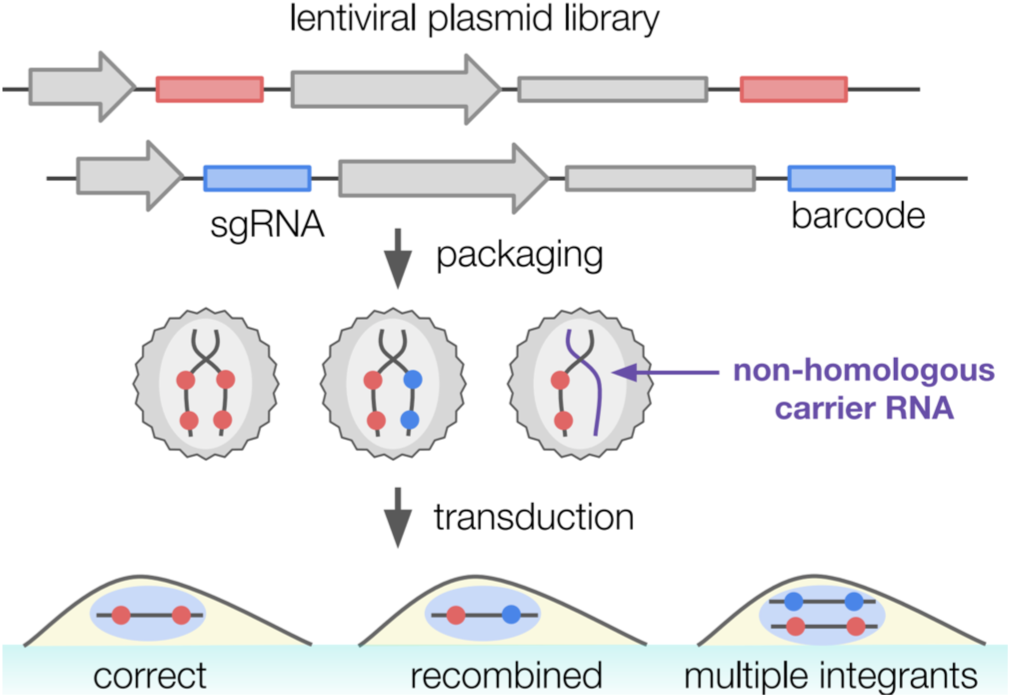
Schematic of delivery of barcoded lentiviral plasmid library into target cells. Viral genomes containing sgRNAs and transcribed RNA barcodes, driven by U6 and EF1a promoters, are packaged as dimers into virions and integrated into target cells. Packaging may yield homodimeric or heterodimeric virions or, in the case of co-packaging with a non-homologous carrier lentivirus (purple), a functionally monomeric virion. Virions with two different library elements have the capacity for recombination between sgRNAs and barcodes as well as potential for integration of multiple perturbations into the target cell, whereas co-packaging with a non-homologous vector limits these alternatives.

## Results

In order to test the feasibility of barcoding a U6-driven sgRNA with a short sequence located in the 3’ UTR of a Pol II-driven resistance transcript, we individually cloned 8 lentiviral plasmids with different sgRNA-barcode pairs and transduced HeLa cells at an MOI < 5%, pooling either before or after lentiviral packaging. After flow sorting and clonally expanding single cells, we analyzed the sgRNA and barcode sequences present in each clone using next-generation sequencing (Methods). We observed that pooled lentiviral packaging resulted in barcode swaps in 37% of clones with a single detected integration, whereas no swapping was detected between individually packaged lentiviral genome sequences (Table 1). Similar results were obtained when packaging a library of 400 barcodes (cloned as a pool). We report barcode swapping rates as the functional and measurable outcome of recombination. Overall, our results are consistent with observations by a number of groups and precautionary comments published in some of the first examples of pooled single-cell gene expression screens^10,11,14^.

**Table 1:** Clonal analysis of lentiviral packaging strategies for barcoded perturbation libraries. Individual transduced cells were isolated by flow sorting and clonally expanded prior to genomic DNA extraction and examination of sgRNA and barcode identity by next-generation sequencing. Each clone was classified by the number of observed integrations, with single integrants further subdivided by whether the sgRNA matched the associated barcode. A- barcode swap rate (column second from right) was estimated by dividing the number of recombined single integrants by the total number of cells with a single integration. Standard packaging refers to packaging of the pooled perturbation library; for arrayed packaging, individual library elements were separately packaged and subsequently pooled for transduction of target cells. For co-packaging approaches, three types of carrier plasmids were used: non-lentiviral (pUC19), integrating lentiviral (pR_H2B-BFP and pLX_TRC313_LacZ) and non-integrating lentiviral (pR_LG). Finally, a quick harvest (11 hours) of the packaged lentivirus was also explored; for all other conditions, virus was packaged for 48 hours.

| packaging condition | # of barcodes | # of cell clones analyzed | single integration, correct association | single integration, barcode swap | multiple integration | estimated barcode swap rate | relative titer |
| --- | --- | --- | --- | --- | --- | --- | --- |
| arrayed | 8 | 48 | 46 (95.8%) | 0 (0.0%) | 2 (4.2%) | 0% | 100% |
| standard | 8 | 61 | 34 (55.7%) | 20 (32.8%) | 7 (11.5%) | 37.0% | 100% |
| standard | 400 | 28 | 16 (57.1%) | 8 (28.6%) | 4 (14.3%) | 33.3% | 100% |
| 10x dilution in pR_LG | 400 | 68 | 67 (98.5%) | 1 (1.5%) | 0 (0.0%) | 1.5% | 1% |
| 1000x dilution in pR_H2B-BFP | 8 | 61 | 59 (96.7%) | 0 (0.0%) | 2 (3.3%) | 0% | 1% |
| 1000x dilution in pLX_TRC313_LacZ | 400 | 53 | 50 (94.3%) | 2 (3.8%) | 1 (1.9%) | 3.8% | 1% |
| 100x dilution in pLX_TRC313_LacZ | 400 | 16 | 12 (75.0%) | 1 (6.2%) | 3 (18.8%) | 7.7% | 3% |
| 1000x dilution in pUC19 | 8 | 45 | 35 (77.8%) | 2 (4.4%) | 8 (17.8%) | 5.4% | 1% |
| standard, quick harvest (11h) | 8 | 51 | 28 (54.9%) | 17 (33.3%) | 6 (11.8%) | 37.8% | 3% |

The standard pooled lentivirus packaging protocol calls for transfecting the packaging cell line with the lentiviral perturbation library and associated packaging plasmids needed to produce virus (Methods). Our clonal analysis of target cells transduced by virus produced in this manner revealed not only barcode swaps but also a higher occurrence of multiple integrants per cell than predicted by a Poisson model of independent integration events, even at MOI below 5%. The presence of multiple genomic integrants even at limiting virus dilutions could be explained by the ability of a single virion to integrate both packaged genomes, or non-independence of integration probability across target cells, possibly as a result of differences in cell state.

Hypothesizing that recombination and multiple integration events are driven by co-packaging of two RNA genomes per lentiviral capsid^13^ and co-delivery of multiple genomes to individual target cells, we tested dilution of the perturbation library in unrelated carrier plasmids as a means of mitigating both of these undesired effects. Three lentiviral plasmids were evaluated as carriers, including two integration-capable vectors (pR_H2B-BFP encoding a histone subunit tagged with blue fluorescent protein, and pLX_TRC313_LacZ, a control vector used in ORF screens). In addition, we tested a non-integrating lentiviral vector with a mutated 5’ long terminal repeat (LTR) and a short LTR-LTR distance of 2.1 kb in hopes of avoiding unnecessary genomic integrations^15^. All three carriers tested reduced recombination rates to 0–4%. Interestingly, we found that while it was necessary to use the integrating carrier plasmids in 1000-fold excess over the perturbation library, the non-integrating carrier plasmid reduced barcode swaps to the same extent at a dilution of only 1:10, perhaps due to enhanced expression of the shorter LTR-LTR transcript in the packaging cell line. Furthermore, co-packaging with a non-homologous lentiviral vector also limited instances of multiple distinct integrations, likely due to a reduction in the probability that two perturbation library variants enter the target cell.

We also tested dilution in a non-lentiviral carrier plasmid, pUC19, hypothesizing that stringent dilution of the perturbation library could reduce the number of library variants in each packaging cell to one or fewer and minimize the risk that heterodimeric virions are produced. This strategy was found to decrease the recombination rate to 6%. However, we still observed 18% of cell clones with greater than one integrated library variant, consistent with the correlated infection hypothesis and indicating that lentiviral plasmids may be a better choice of carrier material.

The main limitation of this dilution strategy is a 100-fold decrease in titer relative to lentivirus prepared with the non diluted perturbation library, measured by counting the number of cell colonies surviving after antibiotic selection. To investigate the trade-off between titer and unwanted lentiviral effects, we titrated the dilution ratio of one of the integrating lentiviral carrier plasmids (pLX_TRC313_LacZ) but found that 100-fold excess did not show the desired performance, with 25% of colonies showing barcode swaps or multiple integrants. Nevertheless, even with diminished viral titer, we have been able to transduce a library of 1,000 perturbations with 300-fold cell coverage.

We also explored whether recombination events hypothetically occurring in the packaging cell line could be reduced by shortening packaging times. We reduced the time from transfection to harvesting viral supernatant from 48 h to 11 h, but observed no decrease in barcode swapping, consistent with a model where most of the recombination occurs in the target cells.

## Discussion

In the context of genetic screens, lentiviral co-packaging can greatly reduce barcode swaps and decrease the background of multiple integrants without constraining library vector design or necessitating individual packaging of library elements. The latter approach was used by Adamson et al. to avoid recombination in a single-cell gene expression screen, but is limited in scalability^14^. Datlinger et al. developed a -specialized CROP-seq vector in which the sgRNA is itself transcribed by Pol II and captured in a 3’ RNA-seq protocol, obviating the need for an additional barcode and eliminating concerns about recombination^5^. However, this approach requires locating the perturbation within the 3’ LTR and is not generalizable to some types of screens (e.g. paired perturbation screens, or screens of regulatory elements monitored via transcribed barcodes)^16^. By addressing both recombination and multiple integration, co-packaged lentiviral libraries have the potential to improve the accuracy of perturbation barcoding and boost the sensitivity of screens that deliver library constructs with multiple variable elements.

We chose to employ clonal analysis of the genomic integrants by next-generation sequencing to achieve sensitive and unbiased detection of perturbation library elements with single-cell resolution. As we amplify each variable sequence in a separate PCR reaction, our approach is not subject to artifacts resulting from PCR-based recombination. Moreover, our readout does not depend on events subsequent to integration and antibiotic selection, such as detection of a fluorescent marker or perturbation of cellular phenotype that may be confounded by multiple integrations. However, clonal analysis is practically limited to the scale of 100–1000 clones per sequencing run, making it better suited for high-confidence measurements of undesired integration events than for systematic optimization across many test conditions.

Under the standard assumption of a zero-truncated Poisson distribution of lentiviral integrations, one would expect a multiple integration rate below 2.5% when transducing cells at an MOI below 5%, However, despite working in this range for our infections, the measured multiple integration rate was greater than 10%, suggesting that, at least in our HeLa cell system, lentiviral integrations detected after antibiotic selection are correlated and do not follow a zero-truncated Poisson distribution as is commonly assumed^17^. It is likely that multiple integration background is a persistent noise source in genetic screens utilizing lentivirus, with an effect that depends on the particular system. A lack of statistical independence between integration events underscores the need to maintain high representation of library elements in transduced cells in order to average over technical and biological noise.

Decreased viral titer is the only drawback we observed to the dilute co-packaging approach. Here, we opted to perform clonal analysis under stringent dilution conditions, which minimized recombination but exacerbated the viral titer issue. For example, using the integration-defective pR_LG 1:10 co-packaging condition, we were able to generate 10,000 HeLa colonies per mL of viral supernatant. At this titer, 35 mL of viral supernatant would be sufficient for one replicate of a screen involving 1,000 perturbations with 300 initially infected cells per library element. The cost of tissue culture and transfection reagents to generate 35 mL of viral supernatant is currently orders of magnitude smaller than the cost of preparing and sequencing single-cell gene expression libraries covering 1,000 perturbations and thus does not pose a limit to the achievable scale of the screen. On the other hand, this titer may be prohibitive for a genome-wide CRISPR screen reading out enrichment of more than 50,000 sgRNAs by amplicon sequencing. Ultimately, any particular screen will exhibit an optimal trade-off between degradation of data quality by lentiviral recombination and loss of titer due to co-packaging with a carrier plasmid, and the user may select an appropriate level of dilution to balance these effects.

The ability to minimize both recombination and multiple integrations by diluting the transfer vector in another lentiviral plasmid supports the hypothesis that RNA dimerization during lentiviral packaging is involved in these undesired outcomes. In the case of dilution with excess lentiviral carrier plasmid, most library genomes are likely packaged with a non-homologous carrier genome such that the frequency of virions containing two library variants is substantially reduced. Meanwhile, limiting dilution with a non-lentiviral carrier plasmid may reduce the likelihood that two library variants are transfected into the same cell for packaging, hence preventing two different library RNA genomes from dimerizing. Both of these approaches mitigate the potential for template switching and recombination between two different library genomes in the target cells.

Alternative approaches to control recombination could include directly inhibiting the template-switching activity of the viral reverse transcriptase in the transduced cells or biasing packaging to a single genome per virion by modifying the RNA sequence or proteins involved in dimerization^18^. Such efforts could potentially address both the effects of recombination and multiple genomic integrations without a corresponding loss in titer due to dilution.

## Methods

Single cell clonal analysis of integrated sgRNAs and barcodes was performed by transducing a lentiviral library into a target cell population, selecting with antibiotic, sorting and expanding single cells, and separately PCR-amplifying and deep sequencing the sgRNA and barcode sequences from each expanded colony. All cell lines were transduced at an MOI <5%, determined by counting the fraction of cells surviving antibiotic selection.

Lentivirus was prepared following published methods^19^. All cell culture used Dulbecco’s Modified Eagle’s Medium supplemented with 10% FBS (GE Life Sciences SH30070.03 T), 100 units/mL penicillin, and 100 μg/mL streptomycin. A 4:3:2 ratio by mass of packaging plasmids pMD2.G (Addgene 12259) and psPAX2 (Addgene 12260), and library transfer vector pLas (Supplementary Sequence 1, lentiviral backbone derived from Addgene 61427) was transfected into 293 FT cells (Thermo Fisher R70007) using Lipofectamine 2000 (Invitrogen 11668019). Fresh media was exchanged 4 h after transfection. At 24 h post-transfection, 2 mM caffeine (Sigma-Aldrich C0750) was added, and at 48 h post-transfection lentiviral supernatant was filtered through 0.45 μm cellulose acetate filters (VWR 28145–481), frozen at -80°C, and thawed immediately before use. HeLa cells (a gift from Dr. Iain Cheeseman’s lab) were infected by mixing lentiviral supernatant with 8 μg/mL polybrene (Sigma-Aldrich 107689-10 G) and centrifuging at 1000 g for 2 h at 33°C. At 6 h post-infection, media was exchanged, and at 24 h post-infection cells were passaged into media containing 300 μg/mL zeocin (Thermo Fisher R25001) and selected for one week. Single cells were sorted into 96-well plates and clonally expanded. More than 90% of wells with cell growth contained single colonies, determined by visual inspection. These expanded clones were analyzed by extracting gDNA, separately amplifying sgRNA and barcode sequences by PCR, and deep sequencing the amplicons (Illumina MiniSeq).

Sequence data from each colony were analyzed by matching reads to known sgRNAs within an allowed edit distance of 2 bases and barcodes within an allowed edit distance of 1 base to accommodate errors in oligo synthesis (Supplementary Table 1). Sequences with fewer than 30 reads or a read fraction below 10% were discarded. Multiple integration events were defined by the presence of more than one sgRNA or more than one barcode sequence. Single integration with barcode swapping was defined as detection of one sgRNA and one barcode cognate to a different sgRNA. We report the frequency of observed barcode swapping events, which does not include multiple integration of the same library element, recombination between two identical library elements, or secondary recombination events that restore the original sgRNA-barcode pairing.

## Acknowledgements

The authors would like to acknowledge Linyi Gao for the gift of plasmid pR_LG, and Linyi Gao, Jonathan Schmid-Burgk, Jacob Borrajo, Mohamad Najia, Atray Dixit, and Charles Fulco for discussions. This work was supported by grant R01 HG009283-01A1 from the National Institute of Health to P.C.B., grant P50 HG006193 from the National Institute of Health to P.C.B, and the Simons Center Seed Grant from MIT with Feng Zhang. Internal funding from the Broad Institute BN10 Photons and BN10 Self Reporting grants also supported this work. P.C.B. was supported by a Career Award at the Scientific Interface from the Burroughs Wellcome Fund.

